# A Minimal Physiological Model of Perceptual Suppression and Breakthrough in Visual Rivalry

**DOI:** 10.1101/2025.05.28.656716

**Authors:** Christopher J. Whyte, Hugh R. Wilson, James M. Shine, David Alais

## Abstract

Visual rivalry paradigms provide a powerful tool for probing the mechanisms of visual awareness and perceptual suppression. While the dynamics and determinants of perceptual switches in visual rivalry have been extensively studied and modeled, recent advances in experimental design - particularly those that quantify the depth and variability of perceptual suppression - have outpaced the development of computational models. Here we extend an existing dynamical model of binocular rivalry to encompass two novel experimental paradigms: a threshold detection variant of binocular rivalry, and tracking continuous flash suppression. Together, these tasks provide complementary measures of the dynamics and magnitude of perceptual suppression. Through numerical simulation we demonstrate that a single mechanism, competitive (hysteretic) inhibition between slowly adapting monocular populations, is sufficient to account for the suppression depth findings across both paradigms. This unified model offers a foundation for the development of a quantitative theory of perceptual suppression in visual rivalry.

## Introduction

When one eye is presented with an image that is different from the other eye’s image, the normal process of binocular fusion is prevented (Blake, 1989, 2001; Blake & Boothroyd, 1985; Fox & Herrmann, 1967). Instead, only one eye’s image is seen while the other image is suppressed from awareness by a process known as interocular suppression. An interesting feature of this paradigm is that the consciously perceived image alternates over time as the two images compete in a kind of ‘rivalry’ for perceptual dominance, with perceptual switches occurring every second or two in a stochastic manner. This process is traditionally studied using binocular rivalry (Fig 1A; Blake, 2001; Blake & Fox, 1974; Levelt, 1965), a paradigm with a long experimental history dating back over a century (Breese, 1899; Creed, 1935) with a rich literature in psychophysics (Alais, 2012; Blake & Logothetis, 2002), neuroimaging (Brascamp et al., 2015; Haynes et al., 2005; Lumer et al., 1998; Polonsky et al., 2000; Tong et al., 1998; Tononi et al., 1998; Weilnhammer et al., 2017) and non-human primate electrophysiology (Dwarakanath et al., 2020; Hesse & Tsao, 2020; Kapoor et al., 2020; Leopold & Logothetis, 1996; Logothetis, 1998; Panagiotaropoulos et al., 2012, 2012; Wilke et al., 2009).

**Figure 1.**
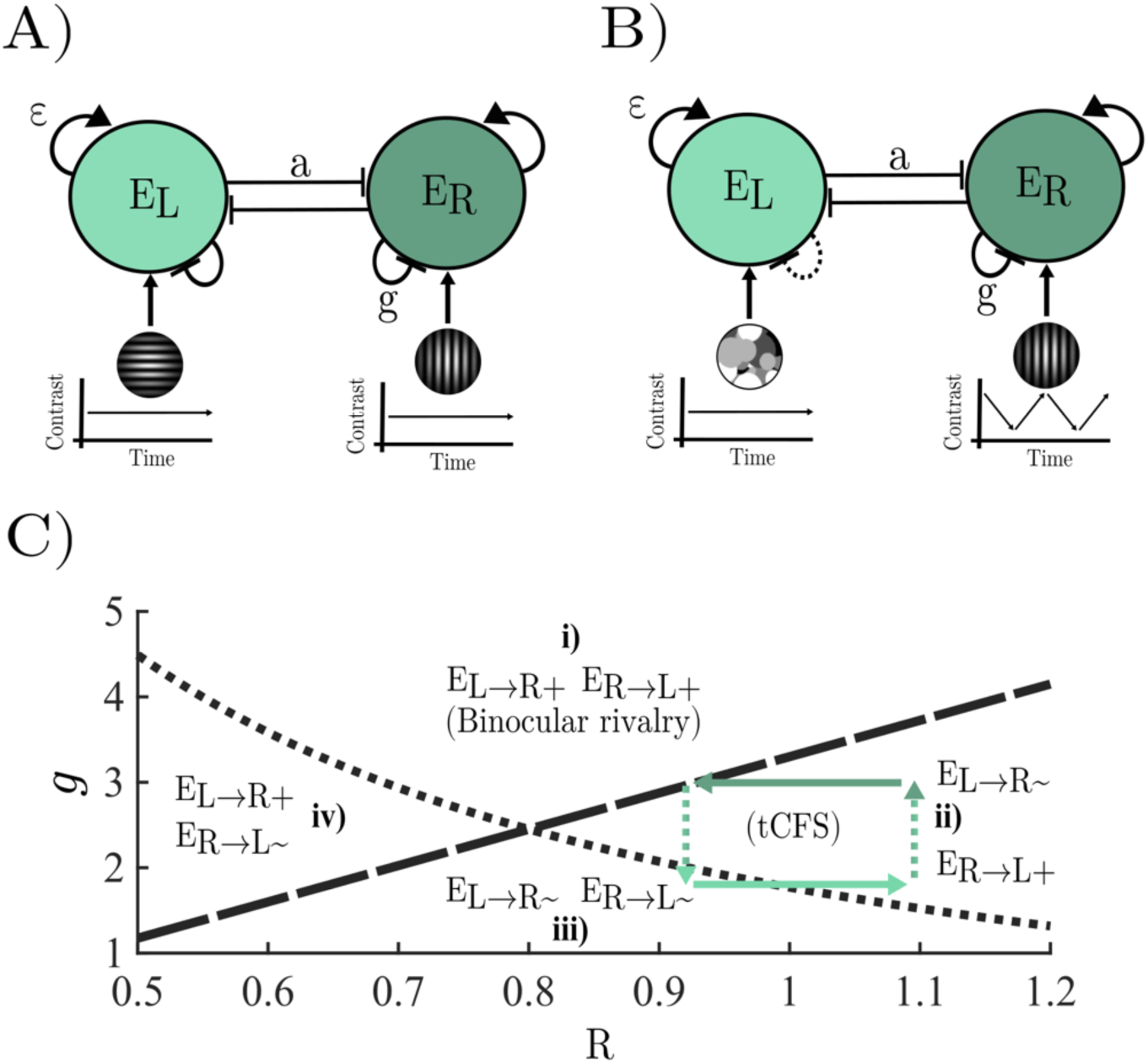
A) Model architecture for binocular rivalry – two monocular populations compete for dominance through a process of mutual winner-take-all inhibition that is asymptotically destabilised by the accumulation of a slow hyperpolarising adaptation current. B) Model architecture for tCFS – two slowly adapting monocular populations compete through mutual inhibition, as they do in binocular rivalry, but instead of being driven by two constant stimuli, the hypothesised effect of the dynamic mask is approximated by driving one population (𝐸_𝐿_) with a stimulus of constant contrast and reducing the strength of adaptation, whilst driving the other population (𝐸_𝑅_)with a dynamic stimulus (equation 6) that increases linearly when the population driven by the target stimulus is suppressed, and decreases linearly when it is dominant. See main text for a more in depth explanation. C) Adaptation by stimulus strength parameter space partitioned into four distinct dynamical regimes by the adaptation strength inequalities in equation 5 (𝑔_𝐿_ − dotted line, 𝑔_𝑅_ − dashed line); regime i) both 𝐸_𝑅_ and 𝐸_𝐿_ are asymptotically released from inhibition by the accumulation of adaptation in the dominant population modelling typical binocular rivalry conditions; regime ii) The population driven by the mask (𝐸_𝐿_) is asymptotically released from inhibition by the target stimulus population (𝐸_𝑅_), but if the mask is dominant adaptation alone will not release the target stimulus population from inhibition; regime iii) both the mask and target stimulus population cannot escape from suppression by adaptation alone; regime iv) the population driven by the mask cannot escape suppression via adaptation alone, whilst the population driven by the target stimulus can escape suppression. Green arrows show a typical trajectory through the parameter space across a full dominance-suppression cycle in tCFS. The target stimulus starts close to full contrast making the strongly adapting target stimulus population dominant (dark green). Stimulus contrast then decreases until the weakly adapting mask driven population (light green) is released from suppression and takes over at which point the contrast of the stimulus starts to increase until the population driven by the stimulus is released from suppression.

Binocular rivalry received a boost in popularity in the 1990s when wide spread interest in the experimental study of consciousness emerged and binocular rivalry was proposed as a key empirical paradigm (Crick, 1995; Logothetis, 1998). The appeal was simple: in rivalry, two visual stimuli enter the visual system but only one is consciously experienced. This offered a practical way to study several interesting questions central to the neuroscience of consciousness. Can a stimulus suppressed from awareness still guide behaviour? Where along the visual pathway does suppression occur? How strong must suppression be to remove a stimulus from awareness? The question concerning rivalry suppression strength has received considerable attention, with many studies attempting to quantify suppression depth(Alais & Melcher, 2007; Apthorp et al., 2009; Blake & Camisa, 1979; Hollins & Bailey, 1981; Lunghi & Alais, 2015; Nguyen et al., 2003; Teng Leng & Loop, 1994; Tsuchiya et al., 2006; Watanabe et al., 2004). The results of these studies show a range of 2–8 dB of suppression, with an average suppression depth of approximately 4-5 dB. Thus, while phenomenal experience in rivalry is that of a stimulus completely disappearing from awareness, measures of suppression depth reveal a relatively modest degree of suppression.

Another form of visual rivalry, known as continuous flash suppression (CFS; Tsuchiya & Koch, 2005), has become very popular in recent years (Gayet et al., 2014; Pournaghdali & Schwartz, 2020; Yang et al., 2014). CFS involves presenting a rapid sequence of high-contrast, random patterns to one eye (the mask) and an image to the other that is usually small, low contrast and static (the target). The mask will reliably suppress the target for much longer periods than is typical in binocular rivalry (for example, 10–20 seconds vs 1–2 seconds) and the dependent variable in CFS is usually the time required for the target to emerge from suppression and reach visual awareness. CFS is claimed to produce stronger suppression than binocular rivalry, although measuring CFS breakthrough times cannot resolve this question. While longer suppression periods in CFS might be due to stronger suppression, it may instead arise from lack of adaptation to the masker due it being continually updated with new random patterns driving unadapted neurons. Due to the fact that binocular rivalry stimuli are sustained, adaptation is thought to reduce suppression depth within each rivalry phase (Alais et al., 2010). To determine CFS suppression strength requires a measure of stimulus contrast thresholds rather than breakthrough times. Separate increment thresholds are needed for breakthrough and suppression so that the visible image’s threshold can be compared with the suppressed image’s threshold, as done in binocular rivalry experiments.

A recent CFS variant known as ‘tracking’ CFS (tCFS; Alais et al., 2023) provides a simple method for measuring CFS suppression depth that enables easy comparisons with estimates from binocular rivalry. It involves a target image that continuously changes in log contrast – either rising of falling – depending on the observer’s perceptual report (Fig 1B). Breakthrough thresholds are registered when a low contrast target that steadily rises in contrast eventually breaks into awareness. As soon as the observer indicates that the target has become visibile, the contrast of the target reverses direction and steadily reduces in contrast until it becomes suppressed once again. As soon as suppression is reported, the target reverses course, and increases in contrast again until breakthrough (and so on, in a cycle). When the cycle ends (e.g., after 10 reversals), the difference between mean thresholds for breakthrough and suppression quantifies suppression depth.

Due to the relative simplicity of binocular rivalry, and the law like relationship between the physical properties of a stimulus and the duration of the resulting percepts (i.e. Levelt’s laws; Levelt, 1965, 1967), formal models of binocular rivalry have been proposed, and iteratively refined and tested, for well over half a century (Brascamp et al., 2015; Lehky, 1988; Levelt, 1965, 1967). This has led to a rich quantitative modelling literature spanning multiple spatial scales and levels of analysis, from implementation level spiking (Laing & Chow, 2002; Wang et al., 2020; Whyte, Müller, et al., 2025; Wilson, 2003) and mean field (Laing et al., 2010; Moreno-Bote et al., 2007; Shpiro et al., 2007, 2009; Wilson, 2007, 2017) models, which describe rivalry dynamics in terms of biophysical processes, to algorithmic and computational level models which cast perceptual switching as optimising Bayesian (Dayan, 1998; Gershman et al., 2009; Gershman et al., 2012; Hohwy et al., 2008; Parr et al., 2019; Whyte et al., 2024) and/or decision theoretic (Safavi & Dayan, 2022, 2024) objectives. Quantitative theory and modelling has, however, lagged behind experimental progress in studying variants of visual rivalry (which we use as a catch all term for paradigms that utilise interocular suppression to render stimuli unconscious; e.g. CFS, and binocular rivalry) that allow researchers to quantify the depth of perceptual suppression (for a notable exception see Shimaoka & Kaneko, 2011).

In this paper we take a first step towards developing a quantitative theory of perceptual suppression in visual rivalry by extending an existing model of binocular rivalry to encompass threshold detection variants of binocular rivalry and tCFS. We begin by introducing the minimal model of binocular rivalry proposed by Wilson (2007). We chose to start from this model as it shares key features common to (almost) all neuronal models of rivalry, namely strong competitive inhibition and self-adaptation (Shpiro et al., 2007). The model also has sufficient complexity to account for the psychophysical “laws” known to govern rivalry (Levelt’s laws; Brascamp et al., 2015; Levelt, 1965)) whilst still being simple enough that it is possible to derive analytic understanding from closed form approximations to the (nonlinear) dynamics of the full network. We use the model to reproduce and explain the results of two experimental paradigms which provide complementary insight into the dynamics of suppression depth in visual rivalry. The first provides an estimate of how suppression depth evolves over time in binocular rivalry (Alais et al., 2010). The second, tCFS (Alais et al., 2023), provides an absolute measure of the magnitude of perceptual suppression by measuring both suppression and breakthrough thresholds.

### Minimal model of visual rivalry

The dynamics of the Wilson (2007) model are governed by two key qualitative features. The first – strong inhibition between competing monocular neuronal populations – generates winner-take-all activity, such that only one monocular population can be active at any one time. The second – strong self-adaptation – guarantees that winner-take-all states are transient, even in the absence of noise, by decreasing the activity of the dominant population sufficiently to release the competing population from suppression via a reduction in competitive inhibition. Together these conditions guarantee a nonlinear oscillation, known as a limit cycle, between two mutually exclusive percepts, separated by brief periods of mixed perception, characteristic of binocular rivalry.

The model is described by a system of four nonlinear ordinary differential equations:

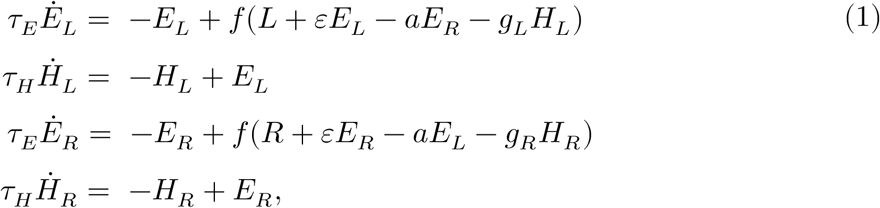

where the population transfer function is the simple threshold linear function 𝑓(𝑥) = max(𝑥, 0), and the four state variables 𝐸_𝐿_, 𝐸_𝑅_ and 𝐻_𝐿_, 𝐻_𝑅_ represent the aggregate neuronal activity of competing monocular populations, and the activity dependent hyperpolarising current of each population, respectively. Each neuronal population is driven by separate monocular input (𝐿, 𝑅), weakly excited (with strength 𝜀) by recurrent projections within each population, strongly inhibited (with strength 𝑎) by the competing population, and undergoes slow self-inhibition (with strength 𝑔) via the accumulation of a slow hyperpolarising adaptation current (𝐻_𝐿_, 𝐻_𝑅_) which represents the action of calcium mediated potassium currents (McCormick & Williamson, 1989; Wilson & Cowan, 2021). In addition, we note that the model assumes that inhibition is purely a function of excitatory activity and rapidly reaches its asymptotic value allowing the potential six-dimensional system of ODEs to be reduced to four.

Working from the qualitative features described above, Wilson (2007) derived a minimal set of sufficient conditions for the presence of a limit cycle in the model. Leveraging the fact that 𝜏_𝐻_ ≫ 𝜏_𝐸_ the dynamics of the model can be partitioned into two phases. In the first instantaneous phase 𝐻_𝐿_, 𝐻_𝑅_ ≈ 0, and in the second asymptotic phase, 𝐻_𝐿_ ≈ 𝐸_𝐿_ and 𝐻_𝑅_ ≈ 𝐸_𝑅_. In both partitions the model can be reduced to a two-dimensional system.

In the first instantaneous partition, if 𝐸_𝐿_, 𝐸_𝑅_ > 0, equation 1 reduces to the following two-dimensional linear system which we write in matrix form:

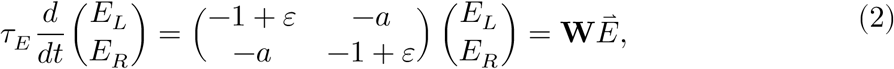

where we have dropped 𝐿 and 𝑅, as external drive shifts the location of the equilibrium (where *Ė*_𝐿_ = *Ė*_𝑅_ = 0), but does not affect its stability (this is equivalent to studying the Jacobian matrix of equation 1). Winner-take-all behaviour requires that the equilibrium of the system is not asymptotically stable, that is, the matrix 𝐖 must have at least one positive eigenvalue. The eigenvalues of 𝐖 are: 𝜆_1_ = 𝜏_𝐸_^−1^(−1 − 𝑎 + 𝜀), and 𝜆_2_ = 𝜏_𝐸_^−1^(−1 + 𝜀 + 𝑎). If 𝑎, 𝜀 > 0, and 𝑎 > 𝜀, which is true for all dynamically plausible parameter values, then 𝜆_1_ < 0, leading to the following inequality for 𝜆_2_ > 0:

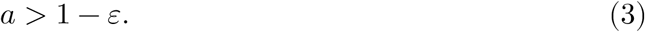

If the inequality in equation 3 is satisfied, unless the system starts in a precisely balanced initial condition with equal external drive (i.e. 𝐸_𝐿_(0) = 𝐸_𝑅_(0) and 𝐿 = 𝑅) the model will converge to a winner-take-all state (i.e. 𝐸_𝐿_ > 0 and 𝐸_𝑅_ = 0 or 𝐸_𝐿_ = 0 and 𝐸_𝑅_ > 0). Once in a winner-take-all state, as long as the competing population is suppressed, the state is stable.

Perceptual switches occur when the suppressed population escapes suppression. For example, if 𝐸_𝐿_ is dominant, and 𝐸_𝑅_ is suppressed, a perceptual switch occurs when 𝐸_𝑅_ escapes from inhibition, which for the simple threshold nonlinearity occurs when the argument of 𝑓(𝑥) passes through zero from below (c.f.(Shpiro et al., 2007)). We, therefore, refer to the argument of the population transfer function as aggregate synaptic drive 𝒟 which we define as a quantity in its own right and show below for the right monocular population:

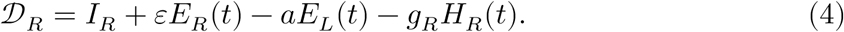

Aggregate synaptic drive takes negative values when the population is suppressed and positive values when dominant. We can find the minimum value of 𝑔_𝐿_ necessary for the right monocular population to escape from suppression asymptotically by demanding that 𝒟_𝑅_ > 0 and solving for 𝑔_𝐿_ leading to:

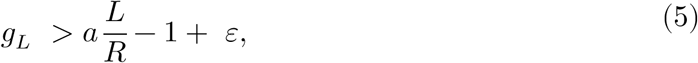

where we have leveraged the fact that when the population is suppressed 𝜀𝐸_𝑅_ = 0, and that at a perceptual switch the model is in the second asymptotic phase of its dynamics where 𝐻_𝑅_ ≈ 0, and 𝐻_𝐿_ ≈ 𝐸_𝐿_. We find the equilibrium value of the dominant population by substituting 𝐻_𝐿_ = 𝐸_𝐿_ into the equation for Ė_𝐿_ = 0, and solve for 𝐸_𝐿_ to obtain 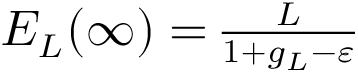 which, when substituted into equation 4, leads to the above inequality. The comparable expression for the asymptotic release of 𝐸_𝐿_ from suppression is given by 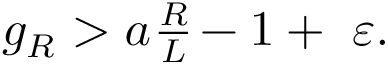 This leaves us with three inequalities that when satisfied guarantee a limit cycle with dynamics that oscillate between mutually exclusive states of perceptual dominance.

To simulate binocular rivalry which presents stationary stimuli with matched image statistics to either eye we set 𝐿 and 𝑅 to constant values with equal adaptation strengths for each monocular population (𝑔_𝐿_ = 𝑔_𝑅_; Fig 1A). Under these conditions (in the absence of noise) the only way for a population to escape from inhibition, and a perceptual switch to occur, is for the model to satisfy the above system of inequalities guaranteeing a limit cycle. These inequalities define the minimal physiological conditions for binocular rivalry in terms of the parameters of the deterministic dynamical system given in equation 1.

Unlike binocular rivalry, where perceptual switches occur every few seconds, in tCFS, perceptual suppression can last for tens of seconds and switches are primarily determined by increasing/decreasing the contrast of the target stimulus. Indeed, in tCFS the stimuli entering both eyes are non-stationary. A dynamic (flashing) mask is presented to one eye and the contrast of the target stimulus presented to the other eye changes linearly with time. We approximated the effect of the dynamic mask (shown to E_L_), which is hypothesised to reduce the accumulation of adaptation, by leaving the value of L fixed and reducing the value of g_L_ relative to g_R_. We arrived at this approximation based upon two considerations. First, although the content of the mask is updated dynamically, the contrast of the mask is constant. Second, in a related modelling study, Shimaoka and Kaneko (2011) showed that the dynamic flashing effect of the mask can be accurately modelled by periodically exciting neurons with different selectivity profiles effectively slowing the accumulation of adaptation. We settled on the precise values of g_L_ and L used in simulation by running a grid search over values of g_L_ and L to find the parameter combination that best minimised the difference between the simulated and empirical dominance durations of Alais et al, (2023) across contrast rates. For full details see (Whyte, Wilson, et al., 2025). The optimal values (g_L_ = 1.7, L = 0.8) violate the inequality in equation 5 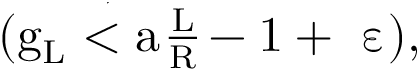 for an adaptation driven release from inhibition (Fig 1B). This means that the population driven by the target stimulus (E_R_) cannot escape suppression by means of adaptation alone.

In tCFS, as described above, the contrast of the target stimulus increases until the population is released from suppression, thereby gaining dominance, and then decreases until it is suppressed again. To model this process we used a piecewise linear function whose value was dependent on which monocular population was dominant:

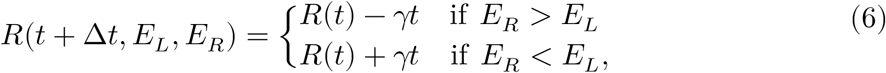

where 𝛾 is a unitless rate constant.

In Fig 1C we show a typical trajectory through the parameter space defined by 𝑔_𝐿/𝑅_ and 𝑅(𝑡) for a single trial of tCFS. The parameter space is partitioned into four regions by the inequalities given by equation 5 which define the model’s dynamical regime. With the exception of a brief period following the suppression of the target stimulus, all values of 𝑅 visited by the model allow the monocular population driven by the mask (𝐸_𝐿_) to asymptotically escape from inhibition via the accumulation of adaptation and do not allow the population driven by the target stimulus (𝐸_𝑅_) to escape from suppression via adaptation alone. That is, escaping from suppression (perceptual breakthrough) requires 𝑅 to increase sufficiently so that 𝒟_𝑅_ > 0, adaptation contributes to this process through a reduction in competitive inhibition from the dominant population, and a reduction in self-inhibition, but is insufficient to lead to a switch without an increase in 𝑅.

All simulations were run by integrating equation 1 numerically using the forward Euler (or Euler-Maruyama when noise was present) method with a time step of 𝑑𝑡 = 0.1 ms in MATLAB 2023b. Model parameters for each simulation are supplied in appendix 1.

### Suppression depth in binocular rivalry

When driven with constant input the model oscillates between mutually exclusive states of perceptual dominance (Fig 2A) defined by the dominance of one monocular neural population and the suppression of the other (Fig 2A upper). Switches occur as the adaptation current approaches its asymptotic value (Fig 2A middle) and the aggregate synaptic drive of the non-dominant population passes through the rectification threshold at zero (Fig 2C lower), allowing the previously suppressed monocular population to inhibit its competitor and gain dominance. Inspection of the aggregate synaptic drive (Fig 2A lower) shows that it has a negative peak shortly after each perceptual switch and then decays exponentially until it passes through the rectification threshold and the suppressed population becomes dominant again.

**Figure 2.**
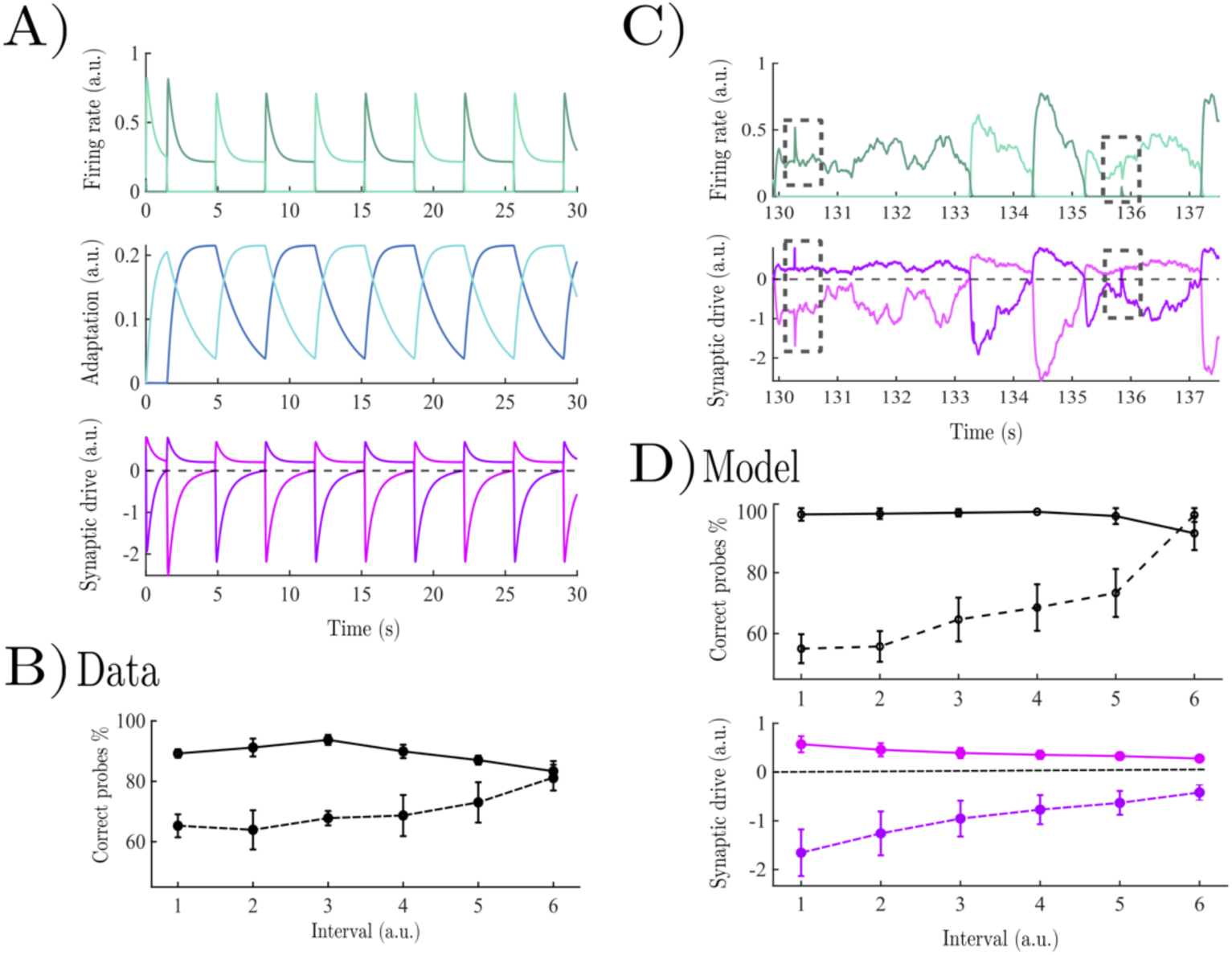
A) Model dynamics for a typical binocular rivalry simulation; upper) firing rate of each competing monocular population; middle) adaptation dynamics; lower aggregate synaptic drive. B) Empirical data adapted from the threshold detection extension of binocular rivalry of Alais et al, (2010). Solid lines show the percentage of correct participant responses to probes presented to the dominant eye, dash lines show the percentage of correct participant responses to probes presented to the suppressed eye. C) Example firing rates and synaptic drive for competing populations receiving probes during periods of dominance and suppression. D) Simulated model behaviour; upper) solid lines show the percentage of correct ideal observer responses to probes presented to the dominant eye, dashed lines show the percentage of correct ideal observer responses to probes presented to the suppressed eye; lower) aggregate synaptic drive for the dominant (solid pink line) population, and suppressed (dashed purple line) populations in each normalised time interval.

This leads to a straightforward prediction, consistent with prior psychological theory (Blake, 1989): the depth of suppression for stimuli with similar selectivity profiles to the rivalry stimuli should decrease as a function of time relative to the last perceptual switch as inhibition from the dominant population, and self-adaptation, decrease. Contrary to this prediction, initial studies found that the detection threshold for stimuli presented to the suppressed eye was constant (Fox & Check, 1972; Norman et al., 2000). However, the stimuli used in each study differed in both spatial frequency and orientation to the rivalry stimuli making it unlikely that the populations excited by the probes were under inhibitory pressure from self-adaptation, or the competing monocular population. In addition, probes were presented in the first half of the suppression period following a perceptual switch where adaptation, which has a time constant on the order of a second (McCormick & Williamson, 1989; Wilson & Cowan, 2021), is unlikely have had sufficient time to exert a large effect on population firing rates. Indeed, in line with the prediction that suppression depth will decrease with time, Alais et al., (2010) found that when probe stimuli matched the spatial frequency and orientation of the rivalry stimuli, and were aligned in time relative to the last perceptual switch, the probability of successful probe detection for stimuli presented to the suppressed eye increased with time relative to the last perceptual switch (Fig 2C dashed line). Similarly, the probability of successful probe detection for stimuli presented to the dominant eye (slightly) reduced with time (Fig 2C solid line).

To understand these results in more detail, beyond the simple qualitative prediction outlined above, we explicitly simulated a threshold detection extension of binocular rivalry of. Following the experimental protocol, each simulation lasted 3 minutes with 60 probes presented to one monocular population at random times over the course of each trial (i.e. a probe was presented approximately every 3 seconds). To generate conditions closer to the *in vivo* experimental conditions we added Gaussian noise (𝜎 = 0.0025) to the adaptation current. For robustness, results were averaged over 40 simulations each with independently generated noise. Each probe was presented for 10 ms of simulation time with strength ranging from [0.1 – 1]. We categorised each probe presentation according to whether the probe was presented in the dominant or suppressed period, and sorted neural activity into 6 time bins normalised by the relative length of each suppression/dominance period. We modelled behavioural responses in terms of what an ideal observer with an optimal criterion (i.e., a criterion selected to minimise misses and false alarms across all six normalised time bins) could readout from the summed population firing rate during the probe presentation window with independent criteria for dominant and suppressed periods (for details see appendix 2). As with the empirical paradigm, probes that induced perceptual switches were excluded from further analysis. For simplicity, in the main text we only show the simulation results for probe strength values of 0.9 but note that the results hold qualitatively across the full range of probe strengths (see appendix 2).

In Fig 2C we show an example of the model dynamics, as a probe is presented to one monocular population in a period of dominance and then suppression. Probes presented whilst the population was dominant generated a spike in the firing rate of the dominant population that could be reliably readout by the ideal observer with occasional misses caused by the noisy adaptation current (solid line Fig 2D upper). Probes presented to the suppressed population broke though less reliably. In Fig 2C the probe coincides with a positive fluctuation in aggregate synaptic drive allowing the aggregate synaptic drive to transiently pass through the rectification threshold and generate a brief spike in the firing rate of the suppressed population. The probability of such breakthroughs occurring increased as a function of time relative to the last perceptual switch (Fig 2D upper). Indeed, the behavioural response curves are essentially a mirror image of the aggregate synaptic drive in each normalised interval flipped about the x-axis (Fig 2D lower).

Here, therefore, in line with empirical data and previous psychological theory (Alais et al., 2010; Blake, 1989), the depth of perceptual suppression in the model decreased as a function of time relative to the last perceptual switch. This is due to the reduction in self-inhibition in the suppressed monocular population, and the reduction in inhibition from the dominant population, both of which were due to the adaptation dynamics pushing the aggregate synaptic drive toward the rectification threshold at zero.

### Suppression depth in tracking continuous flash suppression

The difference in the probability of successful probe detection between the suppressed and dominant eyes in binocular rivalry serves as proxy for the depth of perceptual suppression, an inference supported by the simulations in the previous section where the probability of successful detection mirrors the aggregate synaptic drive, 𝒟. Although having the advantage of being time resolved, this experimental approach does not provide an absolute measure of suppression depth. In this section we generalise the model to tCFS (i.e., ‘tracking’ continuous flash suppression), a paradigm that provides a stationary, but absolute, measure of suppression depth by measuring the threshold for both breakthrough and suppression (Fig 3A; analogous to measuring the difference between the threshold for the successful detection of probes presented to the dominant and suppressed eyes).

**Figure 3.**
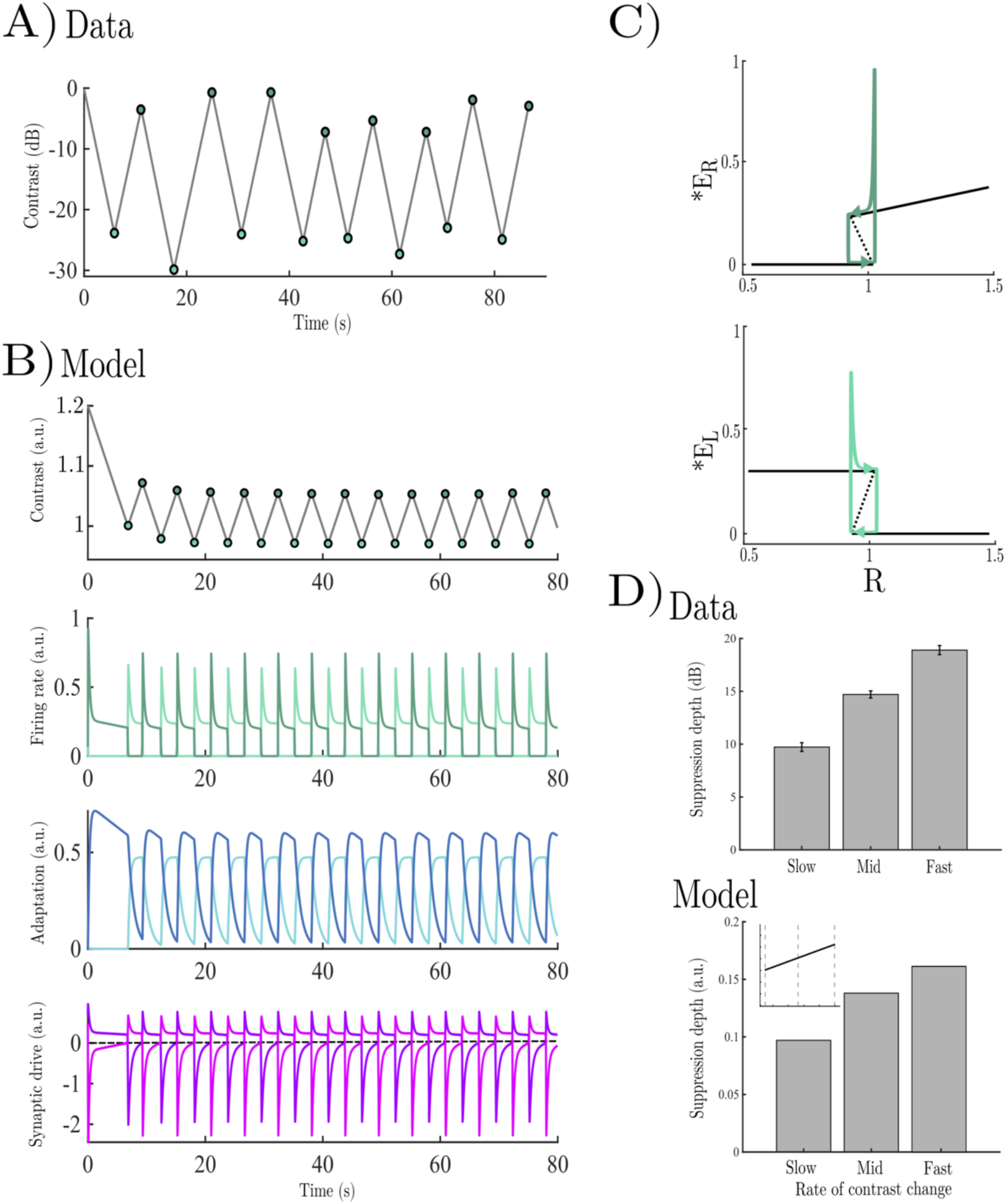
A) Empirical stimulus contrast dynamics from an example tCFS trial adapted from Alais et al, (2023). Light green dots show points of perceptual suppression, and dark green dots show points of perceptual breakthrough. B) Model dynamics for a typical tCFS simulation; upper) stimulus contrast dynamics; upper middle) firing rate of each competing monocular population; lower middle) adaptation dynamics; lower) aggregate synaptic drive. C) Bifurcation diagram for typical firing rate trajectories across the dominance-suppression cycle. Solid lines show the location of stable equilibrium values of *Ė*_𝑅_ = 0 and *Ė*_𝐿_ = 0 when dominant and suppressed. Dashed line shows unstable equilibrium location. D) Suppression depth as a function of contrast rate; upper) empirical data adapted from Alais et al, (2023); lower) simulated data from model, inset shows the three equally spaced values represented bar graph across all simulated contrast rate simulations.

In tCFS, perceptual switches are governed by gradually increasing/decreasing the strength of one monocular stimulus, whilst presenting a high contrast mask to the other eye (Fig 1B). The target stimulus starts at full contrast and then linearly decreases (in decibel scale; dB) until it vanishes from awareness and is suppressed by the mask (indexed by a button press), at which point the stimulus increases in contrast until it escapes suppression from the mask and breaks through into awareness (again indexed by a button press) and the cycle repeats (Alais et al., 2023). Intriguingly, although the threshold for breakthrough varies across stimulus types the difference between breakthrough and suppression thresholds is constant. The threshold for breakthrough is always larger than the threshold for suppression (Fig 3A), suggesting a shared hysteretic mechanism of perceptual breakthrough and suppression that occurs early in visual processing. In addition, the rate of contrast change modulates the depth of perceptual suppression, with faster rates increasing suppression depth (Alais et al, 2023; Fig 3D upper) suggestive of an adaptation driven reduction in suppression depth consistent with the threshold detection binocular rivalry results explored in the previous section (Alais et al, 2010) and the subtractive effect of adaptation in the model.

To formalise and test this hypothesis *in silico*, we simulated tCFS across a range of contrast rates within our minimal model. Importantly, because the mechanism of suppression in the model relies upon competitive inhibition between monocular neural populations, the model is a good representation of the hypothesis that the process of interocular suppression and breakthrough occurs between ocular dominance columns in early visual cortex.

Following the experimental protocol of Alais et al (2023), we simulated tCFS by replacing the constant stimulus driving the right monocular population with a piecewise linear function (equation 6) that linearly decreased/increased the contrast of the target stimulus depending on which monocular population was dominant. We approximated the effect of the dynamic mask by driving the other monocular population with a constant stimulus and reducing the strength of adaptation below the threshold for an adaptation driven release from inhibition based on the hypothesis that refreshing the content of the stimulus reduces the accumulation of adaptation by stimulating neurons with distinct selectivity profiles (Alais et al, 2023).

All simulations started with a full contrast target stimulus which decreased in contrast until the population driven by the mask escaped suppression gaining dominance by inhibiting the stimulus driven population into silence at which point the contrast of the target stimulus increased again and the cycle repeated (Fig 3B). As with the empirical data, the model converged to an equilibrium with stable breakthrough and suppression values within a few contrast reversal cycles post-trial onset, showing the same hysteretic pattern of higher stimulus contrast for breakthrough thresholds than suppression thresholds.

This hysteretic cycle of breakthrough and suppression is shown in the bifurcation diagrams in Fig 3C. Upon breaking free from suppression, each population jumps past the upper-stable branch of the bifurcation diagram and then exponentially approaches the stable steady state given by the expressions for 𝐸_𝑅_(∞) and 𝐸_𝑅_(∞). The population driven by the (decreasing) target stimulus is further from the stable steady state than the mask driven population as the contrast of the stimulus decreases faster than the intrinsic time scale of the system. Instead the system chases, but never reaches, the stable fixed point at 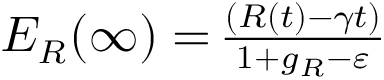 (c.f. Beer, 2022). Perceptual switches occur when the stimulus decreases sufficiently to release the mask from suppression (i.e., when 𝒟_𝐿_ = 0) which then, in turn, suppresses the target stimulus population and the stimulus increases in contrast until 𝒟_𝑅_ = 0. The magnitude of this hysteretic effect (i.e. the difference between the breakthrough and suppression thresholds), is what is quantified by experimental measures of suppression depth. Explaining the determinants of hysteresis in the model, therefore, amounts to explaining the determinants of suppression depth.

The key feature of the model responsible for the presence of hysteresis in the dynamics is the strong recurrent inhibition. This can be seen by deriving approximate expressions for the strength of the target stimulus 𝑅 at breakthrough (𝑅_𝐵_) and suppression (𝑅_𝑆_) points. At a perceptual breakthrough 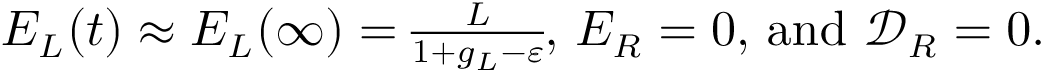 Substituting these values into equation 4 and solving for 𝐼_𝑅𝐵_ we obtain:

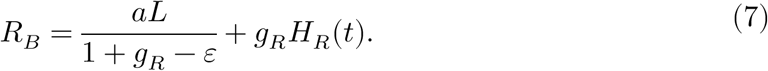

Analogously, when the population driven by the target stimulus is suppressed 𝐸_𝑅_(𝑡) ≈ 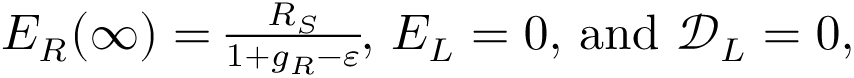 leading to:

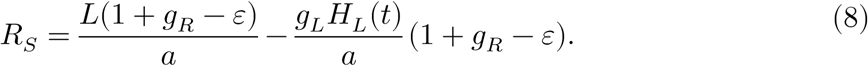

Focusing on the constant terms, we see that competitive inhibition (𝑎) is in the numerator of 𝑅_𝐵_, and in the denominator of 𝑅_𝑆_. This implies that strong competitive inhibition increases the stimulus contrast needed for a population to escape from suppression and decreases the stimulus contrast needed to inhibit the competing population into silence. Whereas increasing the strength of adaptation for the population driven by the target stimulus has the opposite effect.

When the contrast accumulation rate (𝛾) is slow, the contribution of the time dependent adaptation terms is negligible (i.e. 𝐻_𝑅_, 𝐻_𝐿_ ≈ 0), and the magnitude of suppression depth is entirely driven by strong competitive inhibition. As the rate of contrast accumulation increases, adaptation in the suppressed population has less time to decay, increasing suppression depth through the time dependent terms in equations 7-8 which have an additive effect on the breakthrough threshold (𝑅_𝐵_) and a subtractive effect on the suppression threshold (𝑅_𝑆_). This leads to a net increase in suppression depth with increased contrast rate as is seen empirically (𝐅𝐢𝐠 𝟑𝐃). This makes intuitive sense, the subtractive effect of adaptation means that higher external drive from the target stimulus is required to break free from suppression, and likewise, less inhibition is required to keep another population suppressed.

## Discussion

The central appeal of visual rivalry in the experimental study of conscious perception is that two physically-matched stimuli enter the visual system, but only one is consciously experienced, whilst the other is suppressed from awareness. This provides a well-controlled experimental paradigm in which to measure, and potentially mechanistically explain, the threshold for visual awareness and the suppression of content removed from awareness.

In stark contrast to standard binocular rivalry, which has benefited from quantitative neural theory for over half a century, extensions of the standard binocular rivalry paradigm that allow researchers to quantify the depth of perceptual suppression have received little to no attention with the notable exception of Shimaoka and Kaneko (2011) who modified Wilson’s (2007) minimal model to explain the protracted suppression times known to characterise continuous flash suppression (CFS). Building on this work, we extended Wilson’s (2007) model to encompass two empirical paradigms that provide complementary insights into the dynamics (Alais et al., 2010) and magnitude (Alais et al., 2023) of suppression depth in visual rivalry. We used the minimal model of Wilson (2007) as our foundation as it lives in the ‘Goldilocks zone’ of complexity with just enough machinery to account for the key phenomena of interest, whilst still being simple enough that it is possible to derive insights from approximations to the dynamics of the full system. In addition, the dynamics of the model are governed by two key qualitative features, strong competitive inhibition and self-adaptation, common to almost all neuronal models of rivalry (Shpiro et al., 2007). This commonality means that the inferences made in the paper are granted a greater degree of generality than would be possible if we had used a more idiosyncratic model as our starting point.

In the first simulation we showed, in line with empirical data (Alais et al., 2010), that the depth of perceptual suppression in binocular rivalry decreases as a function of time relative to the last perceptual switch. The model shows that this is due to the reduction in self-inhibition in the suppressed monocular population as the population recovers from adaptation accumulated in the preceding dominance period, and a reduction in inhibition from the dominant population as adaptation accumulates. Together, this pushes the aggregate synaptic drive in the suppressed population toward the rectification threshold at zero, increasing the probability with which probe stimuli presented to the suppressed eye will be able to transiently break free from suppression.

In the second simulation, we showed – in line with the predictions of Alais et al (2023) – that the depth of perceptual suppression in tracking continuous flash suppression (tCFS) can be explained by a combination of competitive inhibition and adaptation between monocular populations reminiscent of ocular dominance columns in early visual cortex. When the rate of contrast change is slow, adaptation in the suppressed monocular population has a large amount of time to decay. In this case the hysteretic difference in thresholds for breakthrough and suppression is due solely to competitive inhibition increasing the stimulus contrast needed for a population to escape suppression, and correspondingly, reducing the contrast needed to inhibit the competing population into suppression. As the contrast accumulation rate increases adaptation in the suppressed population has less time to decay and adaptation accumulated in previous periods of dominance carries over across periods of suppression. Crucially, the subtractive effect of adaptation means that suppression depth increases as a function of the rate of contrast change as more external drive from the stimulus is needed in order for the population to break free from suppression, and correspondingly, less inhibition is required to keep the competing population suppressed.

The location of competition in the model most plausibly corresponds to competition between ocular dominance columns within early visual cortex that are biased towards input from one eye (Wilson, 2003, 2007). One possible criticism, therefore, is that the model does not capture findings showing that rivalry can occur at multiple stages in the visual hierarchy depending on stimulus type (e.g. Lee & Blake, 1999; Logothetis et al., 1996). This is an excellent point. Indeed, we are not claiming that rivalry necessarily occur within early visual cortex, rather in line with Wilson (2007), we argue that competitive inhibition between V1-like monocular populations is physiologically sufficient to explain the modulation of suppression depth in standard binocular rivalry and tCFS, a claim that is entirely compatible with multilevel models of more complicated varieties of binocular rivalry which allow competition to occur at a later stage of hierarchical processing (Freeman, 2005; Li et al., 2017; Wilson, 2003). Indeed, our monocular focus is also supported by the current state of the art in non-human primate neurophysiology. In a recent study (Chen et al., 2025) using two-photon calcium imaging it was shown that when monocular-biased neurons in V1 are suppressed by a mask in CFS their orientation tuning amplitude decrease by ∼80%. Another potential criticism is that the model does not capture the finding that suppression depth is constant across stimulus types. Given the simplicity of the model, this is necessarily the case. The model has no spatial extent and, therefore, cannot explain variation in stimulus content. This limitation in scope is, however, common across almost all computational models of binocular rivalry and is a result of the inherent trade off between conceptual tractability and model complexity. As our goal was to explore the minimally sufficient physiological conditions for suppression depth across binocular rivalry and tCFS we leaned heavily on the side of simplicity.

The key contribution of this paper is the unification of threshold detection variants of binocular rivalry and tCFS with one minimally sufficient mechanism. Suppression depth, in both paradigms, can be seen as a result of the same underlying competitive process between slowly adapting monocular populations. In binocular rivalry, the accumulation of adaptation in the dominant population, and the recovery from adaptation in the suppressed population, are the key factors driving the reduction of suppression depth with time. The increase in suppression depth as a function of contrast rate in CFS is a manifestation of this same underlying mechanism. As the contrast rate increases the (subtractive) contribution of adaptation to the aggregate synaptic drive is increased meaning that a larger increase in stimulus contrast is needed for the population to escape inhibition. By the same token, once a population is suppressed, adaptation carried over from the previous dominance period means that less inhibition is required to keep the population suppressed.

Having established the face validity of our model of suppression depth through numerical simulations, in a companion paper (Whyte, Wilson, et al., 2025) we use the same model to generate closed-form expressions for the quantities qualitatively characterised here and move beyond existing experimental data to generate novel empirical predictions based on the model.

## Data availability

Code to reproduce all simulation results reported in the paper can be found at https://github.com/cjwhyte/tCFS.

## Appendix 1: Model parameters

**Table A1.** Parameter description, values, and units of the model described by equations 1-6 for binocular rivalry and tCFS simulations. A longer adaptation time constant was used for the binocular rivalry (BR) simulations to generate empirically realistic dominance durations in the presence of noise.

| Parameter | Description | BR | tCFS |
| --- | --- | --- | --- |
| $\tau_E$ | Neuronal population time constant (ms) | 15 | 15 |
| $\tau_H$ | Hyperpolarizing adaptation current time constant (ms) | 1950 | 1000 |
| $L$ | External drive to left monocular population (a.u.) | 0.85 | 0.8 |
| $R$ | External drive to right monocular population (a.u.) | 0.85 | $R(0) = 1.2$ |
| $\varepsilon$ | Strength of excitatory recurrent projections (a.u.) | 0.05 | 0.05 |
| $a$ | Strength of competitive inhibition (a.u.) | 3.4 | 3.4 |
| $g_L$ | Strength of left population adaptation current (a.u.) | 3 | 1.7 |
| $g_R$ | Strength of right population adaptation current (a.u.) | 3 | 3 |
| $\gamma$ | Rate of contrast change (ms)<sup>-1</sup> | n/a | $(0.0021 - 0.0063) \times 10^{-2}$ |

## Appendix 2: Ideal observer readout

To obtain behavioural responses from the model that are comparable to the participant responses in Alais et al, (2010) we constructed an ideal observer based upon the summed evoked firing rate of the monocular population receiving the probe stimulus. We sorted each probe presentation according to whether they occurred during dominance or suppression periods and sorted neural activity into 6 time bins with normalised durations defined by the length of each suppression and dominance period. As in the empirical analysis, probe presentations that led to a perceptual switch were excluded from further analysis. In addition, we excluded probe presentaions that occurred during dominance and suppressin periods less than 600 ms in duration so that the minimium normalised time window was at least 100 ms. Across probe contrast levels, including probe-absent trials, we calculated the frequency with which the summed firing rate in the 10 ms probe stimulus presentation window exceeded a criterion 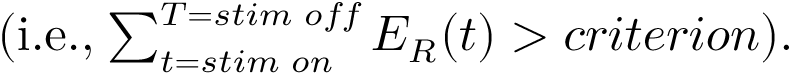 The criterion was defined on the interval between the minium and maximium summed firing rate across all probe strengths. We then selected the (optimal) criterion that best minimised misses and false alarms across all six of the duration-normalised time bins. Criterions for suppressed and dominant populations were calculated independently. Summed firing rates exceeding the optimal criterion were counted as a behavioural response. As the model was not forced to emitt a response we normalised the model response probability (which lie on the inverval [0 – 1]) to vary between [0.5 – 1], under the assumption that in the absence of any information participants (and the model) make an unbiased guess. Model responses across the full range of probe contrast values are shown in Fig A1.

**Fig A1.**
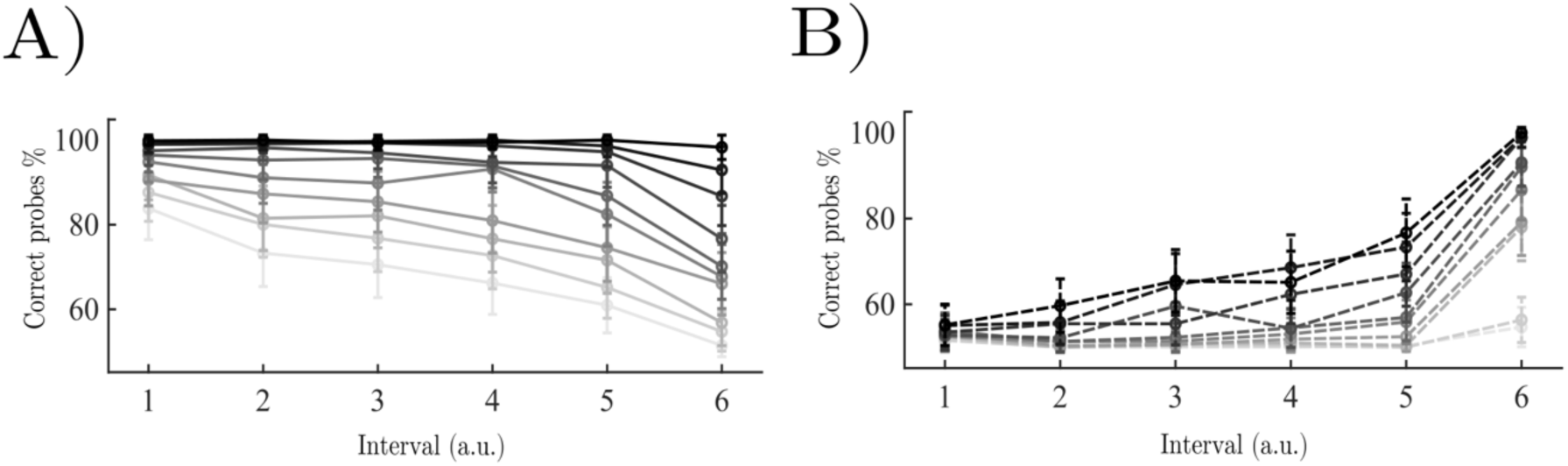
Simulated model behaviour across the full range of probe stimulus contrast values from 0.1 (lightest grey) to 1 (black). A) percentage of correct ideal observer responses to probes presented to the dominant monocular population. B) Same but for the suppressed population.

